# IL-1Ra and CCL5, but not IL-10, are promising targets for treating SMA astrocyte-driven pathology

**DOI:** 10.1101/2023.08.10.552842

**Authors:** Reilly L. Allison, Mya Suneja, Megan LaCroix, Matthew Harmelink, Allison D. Ebert

## Abstract

Spinal muscular atrophy (SMA) is a pediatric genetic disorder characterized by the loss of spinal cord motor neurons. Although the mechanisms underlying motor neuron loss are not clear, current data suggest that glial cells contribute to disease pathology. We have previously found that SMA astrocytes drive microglial activation and motor neuron loss potentially through the upregulation of NFkB-mediated pro-inflammatory cytokines. In this study, we tested the ability of increasing either interleukin 10 (IL-10) or IL-1 receptor antagonist (IL-1ra) while neutralizing C-C motif chemokine ligand 5 (CCL5) to reduce the pro-inflammatory phenotype of SMA astrocytes. While IL-10 was ineffective, IL-1ra ameliorated SMA astrocyte-driven glial activation and motor neuron loss in iPSC-derived cultures *in vitro*. *In vivo* AAV5 delivered IL-1ra overexpression and miR-30 shRNA knockdown of CCL5 had modest but significant improvements on lifespan, weight gain, and motor function of SMNΔ7 mice. Together these data identify IL-1ra and CCL5 as possible therapeutic targets for SMA and highlight the importance of glial-targeted therapeutics for neurodegenerative disease.

## Introduction

5q Spinal muscular atrophy (SMA) was historically a leading genetic cause of infant mortality due to the loss of motor neurons in the spinal cord and the atrophy of skeletal muscle. The advent of novel therapeutic treatments has dramatically impacted patient outcomes, but question remain about the molecular underpinnings of the disease. SMA is caused by variants in the survival of motor neuron 1 (SMN1) gene leading to a significant reduction of SMN protein expression^1^. Although all cell types show reduced SMN protein levels, motor neurons are particularly impacted, and motor neuron loss is the primary phenotypic outcome in SMA patients^2^. SMA motor neurons do show intrinsic deficits in splicing and electrophysiological function; however, simply restoring SMN expression in SMA motor neurons alone does not significantly improve disease pathogenesis^3-6^. These data suggest that other cell types contribute to the vulnerability of motor neurons and overall disease progression in SMA. We have previously shown that astrocytes in SMA are capable of inducing motor neuron loss, have reduced growth factor release, abnormal microRNA production, and increased NFkB expression^7-10^. Others have recently shown that SMA microglia exhibit activation later in the disease process and are involved in synapse engulfment in SMA mice^11^. Using induced pluripotent stem cells (iPSCs) from SMA patients differentiated into astrocytes, microglia, and motor neurons (MNs), our lab has found the ability of SMA astrocyte conditioned media (ACM) to induce SMA microglia activation and motor neuron cell death^10^. These data suggest that aberrantly activated astrocytes drive microglial activation and neurodegeneration in SMA.

Currently, there are three FDA approved treatments for SMA, two of which focus on correcting or adjusting the splicing of SMN2, and the third is an AAV9 gene therapy to deliver functional SMN1. These treatments have proven hugely beneficial to extending patient lifespans and quality of life^12^; however, not all treated patients meet typical developmental milestones, and treatment success is variable across individuals^13-19^. Both clinical data and animal studies indicate that SMN targeted treatments require both early intervention and broad distribution to neurons and astrocytes for best outcomes^20-26^. Based on these data and previous studies showing early astrocyte activation before the onset of motor neuron loss^7^, we hypothesized that targeted treatments to reduce SMA astrocyte pro-inflammatory phenotypes could ameliorate astrocyte-driven microglial activation and neuron loss. Targeted reduction of astrocyte activation in SMA should also allow for both astrocytes and microglia to enact their normal physiological roles in surveillance of the CNS and may allow for an extended therapeutic window in SMA patients.

In this study, we proposed that increasing expression of interleukin 10 (IL-10) – a potent anti-inflammatory ligand produced by astrocytes and microglia in the CNS^27,28^ – specifically in SMA astrocytes would reduce astrocyte malfunction. IL-10 acts via NFkB-dependent and NFkB-independent mechanisms^29-31^, and has been shown to provide direct trophic support to neurons, reduce microglial pro-inflammatory cytokine production^32,33^, and is currently suggested therapeutic target in multiple diseases^34^. We found that treatment with IL-10, even when combined with neutralization of upregulated pro-inflammatory C-C motif chemokine ligand 5 (CCL5/ RANTES), was ineffective at reducing SMA astrocyte activation phenotypes, preventing SMA microglial phagocytosis, or improving SMA MN function. We instead discovered that increasing IL-1ra, a natural antagonist of the IL-1 receptor^35^ produced by astrocytes and microglia in the CNS^36^, in combination with CCL5 knockdown was a more appropriate treatment for reducing astrocyte activation and astrocyte-driven pathology in both mono- and co-cultures of SMA iPSC-derived microglia and MNs *in vitro*. Finally, we used an AAV-based gene therapy approach in the SMNΔ7 mouse model and found a small but significant improvement of lifespan, motor function, and weight gain associated with retained SMI-32+ neurons in the lumbar spinal cord following simultaneous IL-1ra upregulation and CCL5 knockdown in treated mice. Together, these data indicate that astrocyte-mediated inflammatory signaling impacts SMA disease phenotypes *in vitro* and *in vivo* and may provide novel targets for future therapeutic intervention.

## Results

SMA patient-derived induced pluripotent stem cells (SMA iPSCs) were differentiated into spinal-cord patterned GFAP-positive astrocytes (Figure 1A) using established protocols^10,37,38^. To identify a secretome profile specific to SMA astrocytes, SMA astrocyte conditioned media (ACM) was examined via multiplex human cytokine array and compared to healthy control (HC) ACM for dysregulated cytokine secretions (Figure 1B). SMA ACM was found to contain approximately 25-fold higher secretions of C-C motif chemokine ligand 5 (CCL5/ RANTES) than HC ACM. Astrocytic CCL5 release has been shown to increase microglial chemotaxis and activation through CCR1, CCR3, and CCR5 binding^39^ and could contribute to SMA astrocyte-driven pathology (Figure 1C). Consistent with previous studies, SMA iPSC-differentiated microglia exposed to SMA ACM for 24 hours increased priming and activation phenotypes via live imaging analyses (Figure 1D) demonstrated through a significant increase in soma size compared to HC ACM (Figure 1E, 1-way ANOVA, HC ACM vs SMA ACM ****p<0.0001) and a significant increase in SMA microglial phagocytosis of pH-indicator beads with SMA ACM treatment compared to untreated (UTX) or HC ACM treatment (Figure 1F, 1-way ANOVA, UTX vs SMA ACM ***p=0.0002, HC ACM vs SMA ACM ***p=0.0001). Dysfunction of SMA iPSC-differentiated motor neurons (MNs, Figure 1G) has been observed^8,40,41^, including deficits in firing^38,42,43^ which are exacerbated by SMA astrocytes^38^. SMA MNs exhibit increased calcium activity in response to the depolarizing stimuli KCl and glutamate (Figure 1H, 2-way ANOVA, ****p<0.0001). Calcium flux of SMA MNs increases with HC ACM treatment but decreases with SMA ACM treatment (Figure 1I, 1-way ANOVA, *p=0.0275, **p=0.0021). This indicates the ability of HC astrocyte secreted factors to support SMA neuronal activity that are potentially lost in aberrantly activated SMA astrocytes.

**Figure 1.**
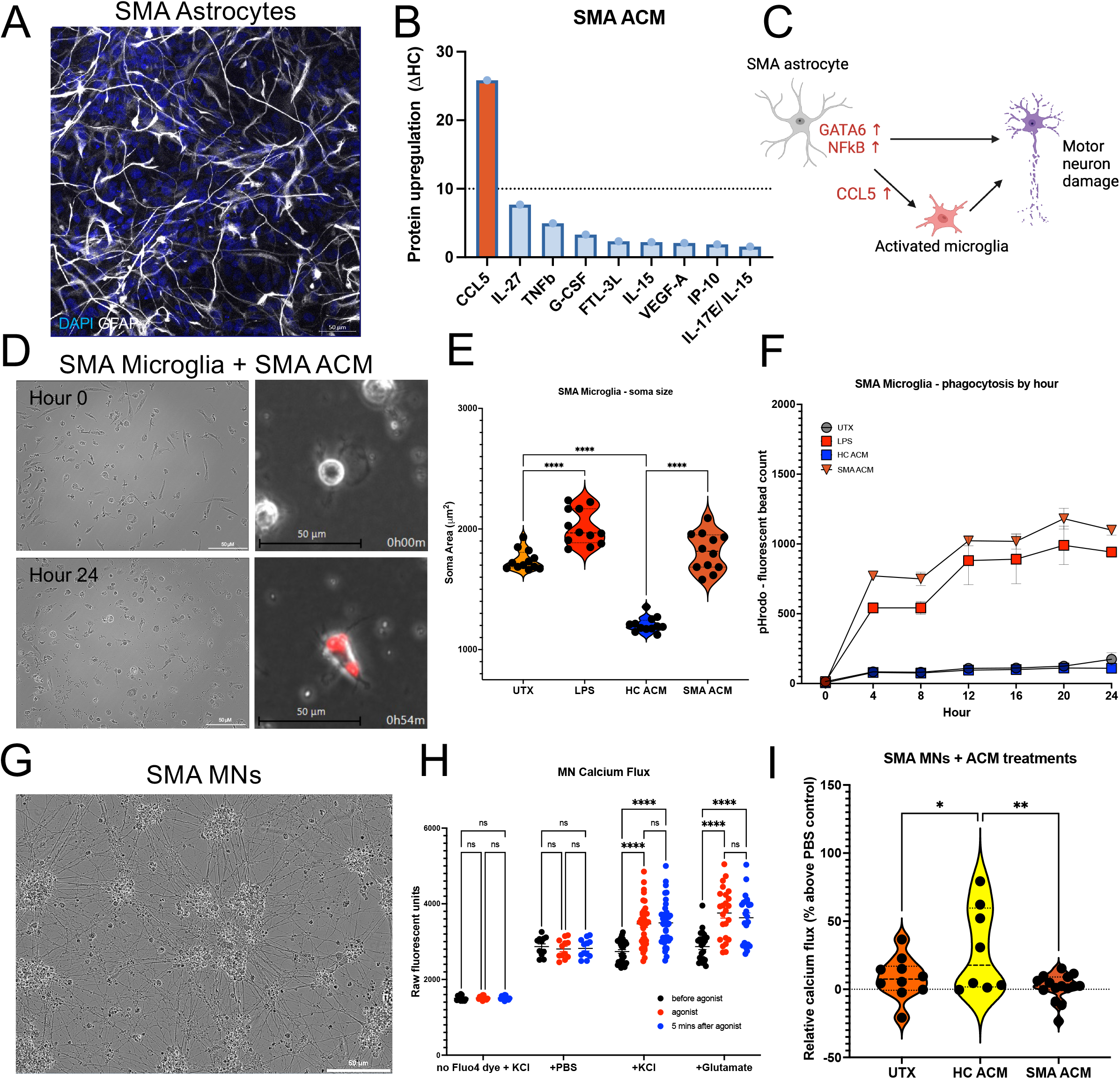
SMA astrocytes secrete high levels of pro-inflammatory factors, drive microglial phagocytosis, and induce MN dysfunction. Astrocytes differentiated from SMA patient iPSCs (**A**, representative image, 63x objective) show aberrant activation. **B.** Multiplex human cytokine array on astrocyte conditioned media (ACM) reveals 25-fold higher secretions of pro-inflammatory cytokine CCL5 compared to healthy control (HC) iPSC-derived astrocytes. **C**. Working theory of astrocyte-driven microglial activation and MN damage involving NFkB-mediated CCL5 secretions. **D**. Representative images of SMA iPSC-derived microglia increase in soma size (left, 20x objective) and phagocytosis of pH-indicator beads (right, 20x objective) after exposure to SMA ACM. SMA microglia show a significant increase in soma size (**E**) and phagocytosis (**F**) after exposure to LPS or SMA ACM compared to SMA microglia treated with HC ACM (1-way ANOVAs, ****p<0.0001). SMA iPSC-derived MNs (**G**, representative image, 20x objective) show increased calcium flux in response to depolarization with KCl and glutamate (**H**, 2-way ANOVA, ****p<0.0001) which significantly increases after exposure to HC ACM and decreases after exposure to SMA ACM (**I**, 1-way ANOVA, *p<0.05, **p<0.005).

To reduce the pro-inflammatory activation phenotype of SMA astrocytes and SMA astrocyte-driven pathology, we treated SMA astrocytes with 430pg/mL of recombinant human interleukin 10 (IL-10), a potent anti-inflammatory ligand that acts to reduce inflammation via inhibition of NFkB and through other NFkB-independent mechanisms^29-31^, and provide direct trophic support to neurons and reduce microglial pro-inflammatory cytokine production^32,33^. To ensure reduction of SMA astrocyte-driven microglial activation and prevent autonomic pro-inflammatory signaling via CCL5, we combined IL-10 treatment with 20pg/mL of CCL5 neutralizing antibodies (NAb) (IL10/CCL5NAb, Figure 2A). We also included a 48-hour wash condition as a control for lingering IL-10 or CCL5 NAb and to assess the impacts of a non-continuous treatment on astrocyte phenotypes (Figure 2B). We did not find any significant improvements of IL10/CCL5NAb on SMA astrocyte transcript production for NFkB or downstream pro-inflammatory ligands IL-1β or IL-6, which are known to be upregulated by SMA astrocytes^7-10^ (Figure 2C, 2-way ANOVAs, ns). We did see a significant reduction in transcripts for complement factors C1q and C3, though transcripts increased when treatments were removed in 48hr wash condition (Figure 2C, 2-way ANOVAs, *p=0.0438, ****p<0.0001). We also did not find any beneficial increases in either glial-derived neurotrophic factor (BDNF) or brain-derived neurotrophic factor (BDNF) transcript production (Figure 2C, 2-way ANOVA, *p=0.0268, ns). Despite expressing the proteins necessary for IL-10 signaling (Figure 2D), SMA astrocytes did not increase expression of IL-10 nor downstream targets of IL-10 receptor activity^27,44,45^ including suppressor of cytokine signaling 3 (SOCS3) or phosphorylated signal transducer and activator of transcription 3 (phSTAT3) after IL10/CCL5NAb (Figure 2E, 1-way ANOVAs, ns). There was a trend for increased phosphorylated cAMP response binding protein (pCREB) with IL10/CCL5NAb and a significant decrease when treatment was removed in 48hr wash condition (1-way ANOVA, UTX to IL10/CCL5NAb p=0.0907, IL10/CCL5NAb to 48hr wash *p=0.0402). pCREB is inhibited by CCR5 activity^39,46,47^, so this increase may indicate a benefit of CCL5 neutralization on pro-inflammatory autocrine signaling in SMA astrocytes rather than an effect of the IL-10 treatment. Application of SMA IL10/CCL5NAb ACM onto SMA microglia did reduce the soma size priming phenotype, although this effect was lost in the 48hr wash condition (Figure 2F, 1-way ANOVA, SMA ACM to SMA IL10/CCL5NAb ACM ****p<0.0001, SMA IL10/CCL5Nab to SMA 48hr wash ACM ****p<0.0001). This reduction in priming was associated with a decrease in phagocytosis, although again the effect was lost when treatment was removed (Figure 2G, 1-way ANOVA, SMA ACM vs SMA IL10/CCL5NAb *p=0.0446, SMA ACM vs SMA 48wash ACM p=0.9571). Direct treatment of SMA microglia with IL-10 and CCL5 NAbs did not reduce the negative impact of SMA ACM treatment (Figure 2G, 1-way ANOVA, ns). Application of SMA IL10/CCL5NAb ACM also did not increase calcium flux in SMA MNs (Figure 2H, 1-way ANOVA, ns). These findings indicate a potential benefit to CCL5 neutralization on SMA astrocyte-driven microglial activation but suggest that IL-10 treatment has a limited impact on SMA astrocyte activation phenotype or astrocyte-driven pathology.

**Figure 2.**
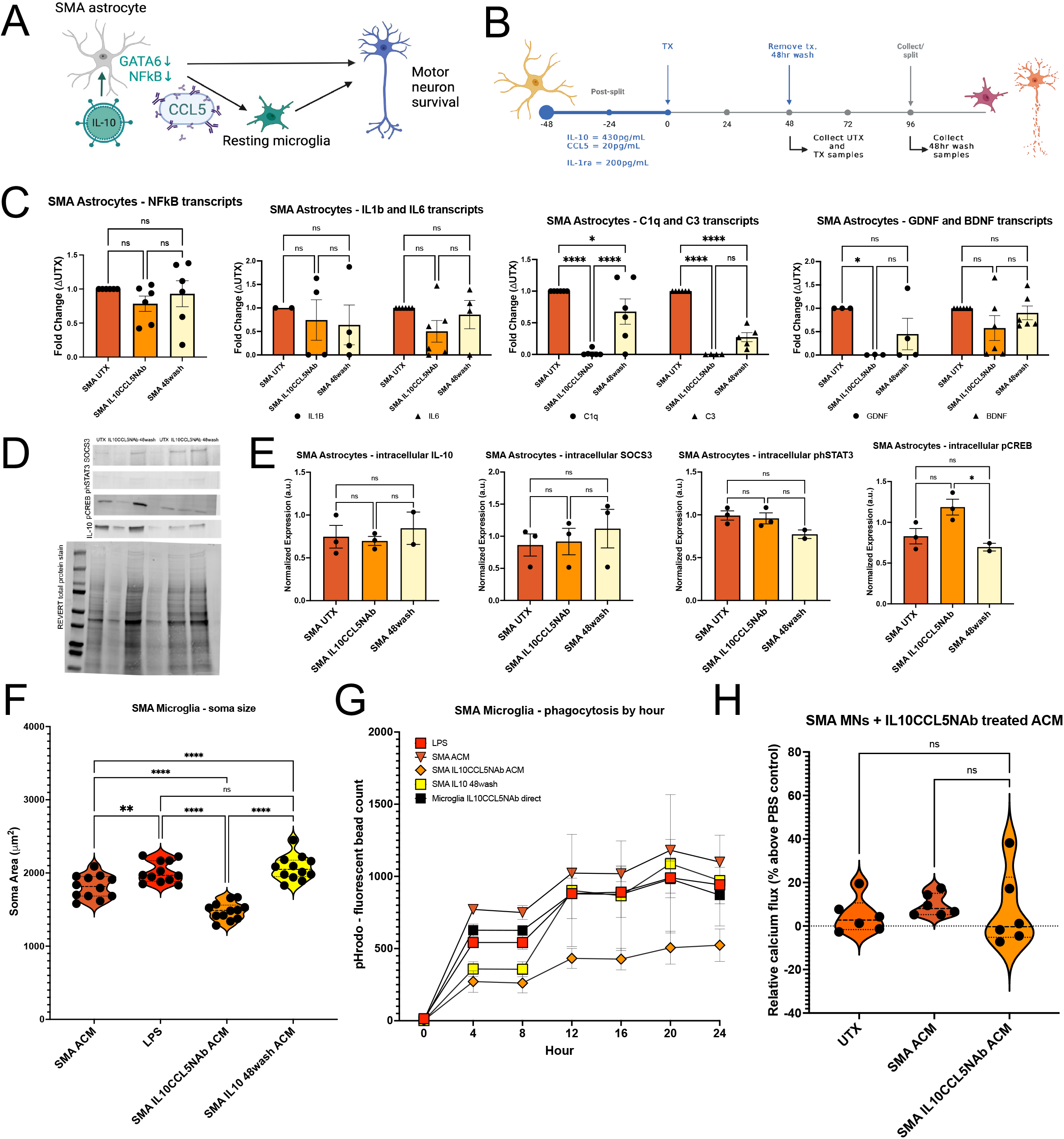
Treating SMA astrocytes with IL-10 and neutralizing CCL5 is not sufficient to reduce activation phenotype. To reduce activation and pro-inflammatory ligand production, SMA astrocytes were treated for 48 hours with 430pg/mL of anti-inflammatory ligand IL-10 and 20pg/mL of CCL5 neutralizing antibodies (**A**, schematic and **B**, treatment paradigm). A 48-hour wash condition was included to assess impacts after treatment was removed (48wash). **C.** No decreases in transcripts for NFkB or downstream pro-inflammatory ligands IL-1β or IL-6 were found after treatment (IL10/CCL5NAb) compared to untreated (UTX) astrocytes. IL10/CCL5NAb decreased transcripts for complement factors C1q and C3, though transcript increase was observed after treatment was removed from astrocytes (48wash). No increases in neurotrophic factor transcript production were found (1-way ANOVAs, ns, *p<0.05, ****p<0.0001). **D**. Western blot analyses for IL-10 signaling reveals no significant increases in downstream IL-10 signaling targets phSTAT3, SOCS3, or increases in IL-10 (**E**, 1-way ANOVAs, ns). Decreased pCREB, a downstream target inhibited by CCR5 signaling, was observed after removal of treatment indicating a possible effect of CCL5 neutralization (1-way ANOVA, *p<0.05). **F.** Treatment of SMA microglia with IL10/CCL5NAb ACM decreases soma size compared to SMA UTX ACM, though this effect is lost in 48wash ACM condition (1-way ANOVA, **p<0.005, ****p<0.0001). **G**. No significant decrease in phagocytosis of pH-indicator beads was found after IL10/CCL5NAb ACM or 48wash ACM treatment (1-way ANOVA, ns). **H**. SMA IL10/CCL5NAb ACM also did not significantly increase MN calcium flux compared to untreated or SMA UTX ACM treated SMA MNs (1-way ANOVA, ns).

Interestingly, we did find that IL10/CCL5NAb increased SMA astrocyte secretions of the anti-inflammatory ligand interleukin 1 receptor antagonist (IL-1ra) when a second IL10CCL5 treatment was applied in place of the 48hr wash (96hrtx) (Figure 3A, 1-way ANOVA, *p=0.0480). IL-1ra is a direct antagonist to the IL-1 receptor and functions by blocking the binding of IL-1a and IL-1β35 to prevent downstream activation of NFkB (Figure 3B). Because IL-1β is upregulated by SMA astrocytes *in vitro^10^* and *in vivo^9^*, we adapted our treatment paradigm to replace IL-10 with IL-1ra as the anti-inflammatory treatment coupled with CCL5 NAbs (IL1ra/CCL5NAb, Figure 3C). SMA IL1ra/CCL5NAb astrocytes had significantly reduced transcripts for NFkB and a strong trend for this maintained reduction even after treatment was removed (Figure 3D, 1-way ANOVAs, UTX to IL1ra/CCL5NAb *p=0.0137, UTX to 48hr wash p=0.0606). These changes were associated with a slightly delayed reduction of transcripts for IL-1β and IL-6 in the 48hr wash condition (Figure 3D, 1-way ANOVAs, *p=0.0404, **p=0.0047). C1q transcripts were again significantly reduced in the SMA IL1ra/CCL5NAb condition compared to UTX (Figure 3D, 1-way ANOVAs *p=0.0123), whereas C3 transcripts were significantly reduced with IL1ra/CCL5NAb (**p=0.0031) and had a nearly significant reduction in the 48hr wash condition (p=0.0645). Despite decreased pro-inflammatory transcripts, there was still a reduction in GDNF and BDNF transcript expression (3D, 1-way ANOVAs, *p=0.0114). Western blot analyses of protein expression found NFkB significantly decreased in SMA IL1ra/CCL5NAb and both NFkB and phNFkB decreased in 48wash condition compared to SMA UTX astrocytes (Figure 3E, 3F, 1-way ANOVAs, NFkB UTX vs IL1ra **p=0.0027, UTX to 48wash ***p=0.0003, phNFkB UTX vs IL1ra p=0.0992, UTX to 48wash *p=0.0164). pCREB protein expression was significantly higher in SMA IL1ra/CCL5NAb astrocytes compared to UTX again indicating a successful reduction in autocrine CCR5 signaling with IL1ra/CCL5NAb (Figure 3F, 1-way ANOVA, *p=0.0246). Compared to SMA UTX ACM, application of both SMA IL1ra/CCL5NAb and 48wash ACM onto SMA microglia (Figure 3G) significantly reduced both microglia soma size (Figure 3H, 1-way ANOVA, ****p<0.0001) and phagocytosis (Figure 3I, 1-way ANOVA, UTX to IL1ra/CCL5NAb ***p=0.0003, UTX to 48wash **p=0.0015). Importantly, direct treatment of SMA microglia with IL-1ra and CCL5 NAbs did not reduce the effect of SMA ACM on these cultures (Figure 3I, ns). These findings reveal the necessity for astrocyte-targeted treatments to reduce disease pathology and pro-inflammatory phenotypes.

**Figure 3.**
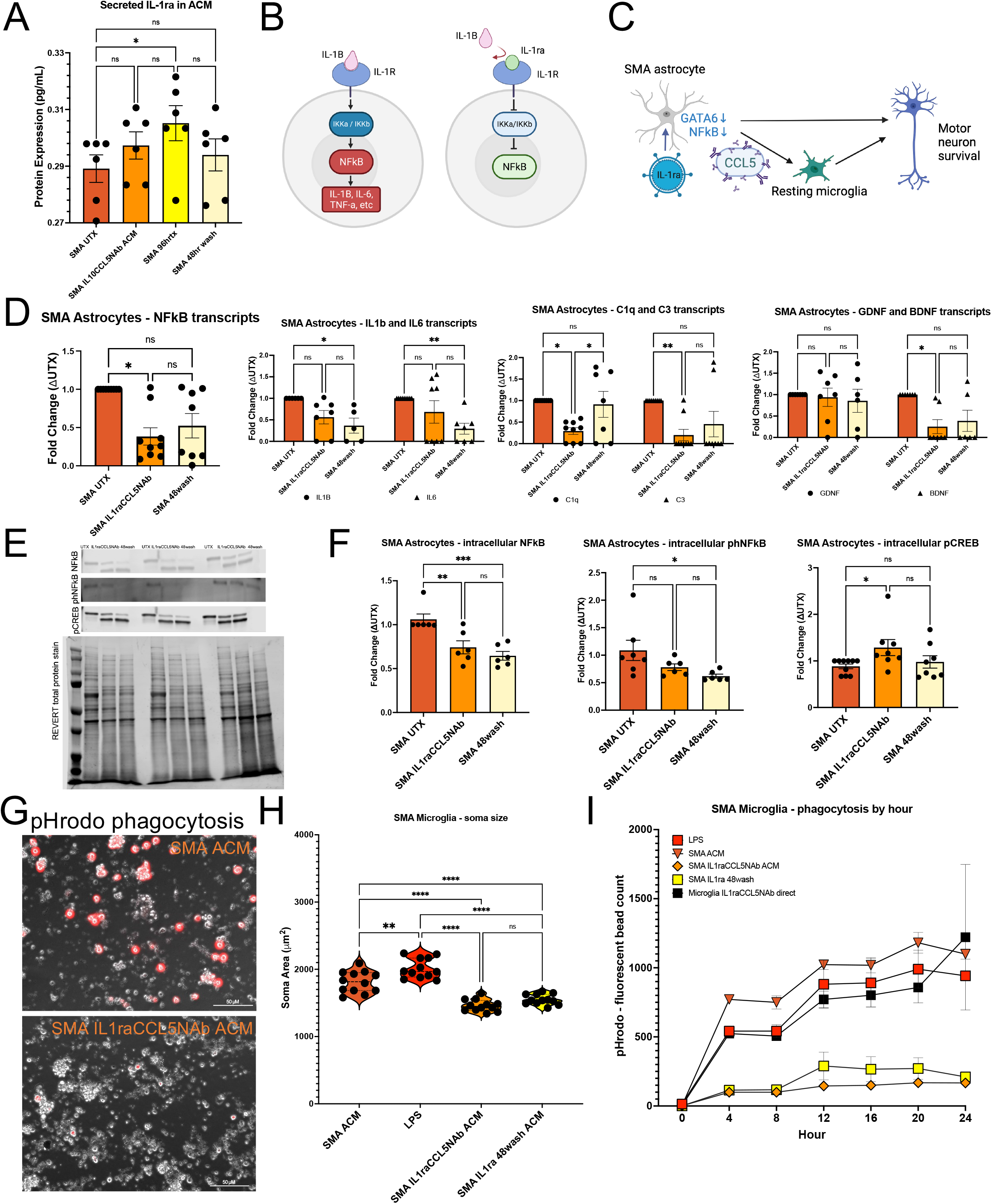
Inhibition of IL-1 signaling through IL-1Ra treatment is more specific and appropriate to reduce SMA astrocyte activation and pathology. **A**. SMA IL10/CCL5NAb astrocytes increased production of IL-1ra after 96-hour sustained treatment (1-way ANOVA, *p<0.05). IL-1ra can reduce NFkB-mediated pro-inflammatory ligand production through inhibition of IL-1 signaling (**B**, schematic and **C**, new treatment paradigm with IL-1ra in place of IL-10). **D**. Treatment of SMA astrocytes with 200pg/mL IL-1ra and 20pg/mL of CCL5 neutralizing antibodies (IL1ra/CCL5NAb) significantly decreased transcript production for NFkB which was associated with a decrease in transcripts for IL-1β and IL-6 48-hours after treatment was removed (1-way ANOVAs, *p<0.05, **p<0.005). Decreased C3 and C1q transcripts were maintained in new treatment paradigm and again there were no increases in transcripts for neurotrophic factors (1-way ANOVAs, ns). **E**. Western blot analyses confirm decreased NFkB protein expression during treatment and maintain this effect 48-hours after treatment removal (**F**, 1-way ANOVA, **p<0.005, ***p<0.0005). Phosphorylated NFkB protein was also decreased in the 48wash condition compared to UTX (1-way ANOVA, *p<0.05). pCREB protein was increased in IL1ra/CCL5NAb astrocytes though this increase was not significant in 48wash condition (1-way ANOVA, *p<0.05). Treatment of SMA microglia with IL1ra/CCL5NAb ACM or 48wash ACM (**G**, representative image) significantly decreased priming (**H**) and phagocytosis (**I**) (1-way ANOVAs, **p<0.005, ****p<0.0001).

We next examined the effects of SMA IL1ra/CCL5NAb astrocytes on co-cultures of SMA microglia and SMA motor neurons. Lentiviruses driving expression of either GFP or RFP under the elongation factor 1 (EF-1) promotor were used to establish stable SMA patient lines of fluorescent iPSCs which could be differentiated into microglia and MNs (Figure 4A). SMA microglia did not have measurable calcium flux in response to KCl in high-throughput calcium assay (Figure 4B, 2-way ANOVA, ns), nor did the addition of SMA microglia significantly alter the calcium flux of SMA MNs without the influence of astrocytes (Figure 4C, t-test, ns). With the addition of SMA IL1ra/CCL5NAb ACM, SMA MNs co-cultured with SMA microglia had significantly increased calcium flux (Figure 4D, 1-way ANOVA, SMA ACM to IL1ra/CCL5Nab *p=0.0222). Phagocytosis in co-cultures was assessed in live imaging assay as fluorescent microglia became positive for both GFP and RFP signal upon phagocytosis of oppositely labeled fluorescent MN (Figure 4E). After 48-hours of co-culture treatment with ACM, phagocytic GFP+/RFP+ microglia were significantly reduced in both IL1ra/CCL5NAb and 48wash ACM conditions (Figure 4F, SMA UTX ACM vs IL1ra/CCL5NAb ****p<0.0001, SMA UTX ACM vs 48wash ****p<0.0001). Again, there was no significant reduction when co-cultures were treated directly with IL-1ra and CCL5 NAbs with SMA UTX ACM vs SMA UTX ACM alone (Figure 4F, 1-way ANOVA, ns). Co-cultures were next assessed for MN apoptosis via TUNEL assay (Figure 4G, Supplemental Figure S2). SMA IL1ra/CCL5NAb ACM significantly reduced SMA MN apoptosis compared to both untreated and SMA ACM treated co-cultures (Figure 4H, 1-way ANOVA, UTX vs IL1ra/CCL5NAb *p= 0.0161, SMA ACM vs IL1ra/CCL5NAb ****p<0.0001). These co-culture data suggest the ability of SMA astrocyte-targeted IL-1ra and CCL5 treatments to reduce glial inflammatory phenotypes and disease pathology.

**Figure 4.**
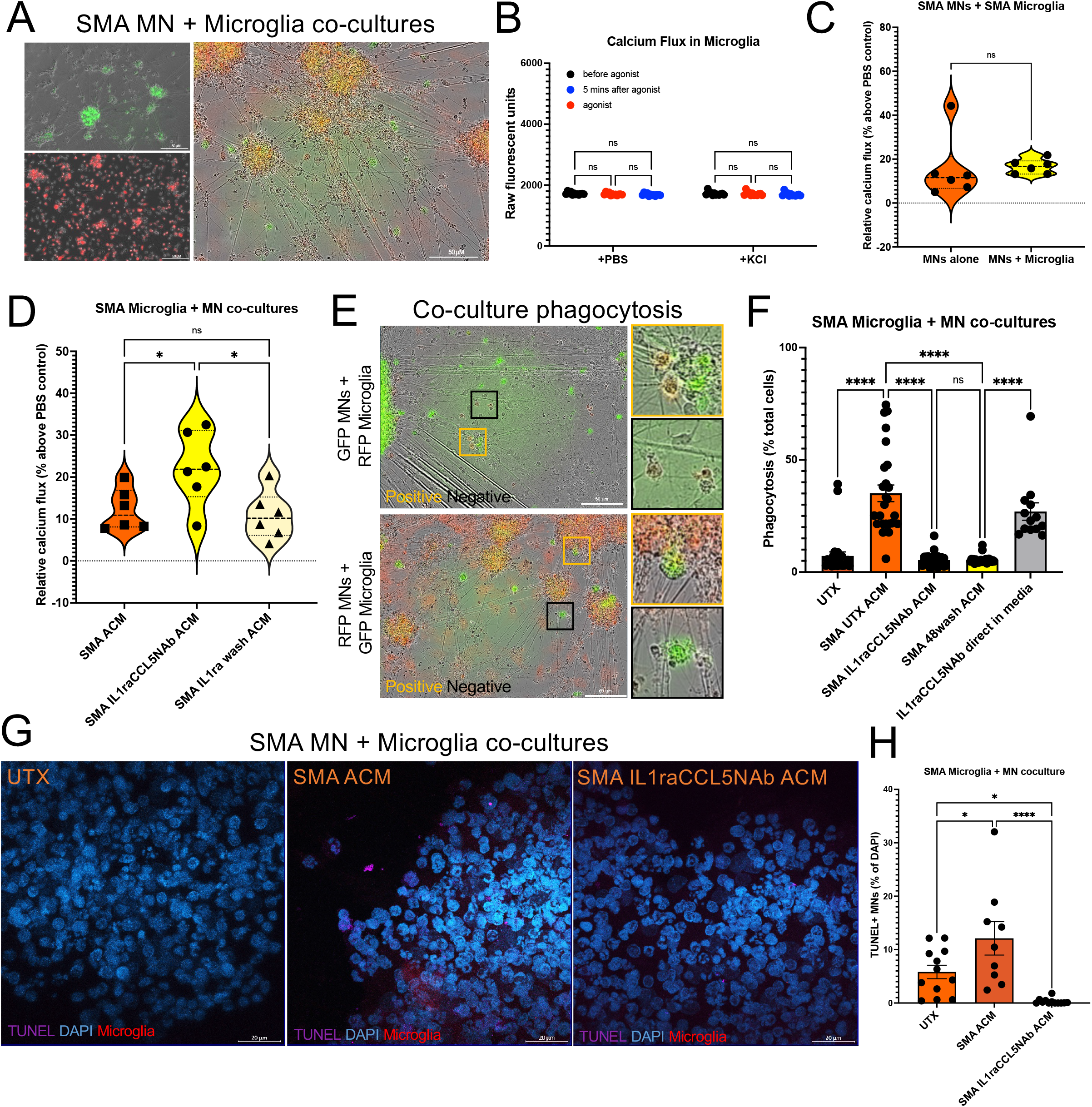
Co-cultures of SMA microglia and SMA MNs have decreased phagocytosis and MN loss when exposed to SMA IL1ra/CCL5NAb ACM compared to SMA UTX ACM. **A**. Stable lines of GFP and RFP SMA iPSCs differentiated into MNs (left, top) and microglia (left, bottom) to allow for live imaging of co-cultures (right). **B**. Microglia alone do not have measurable calcium flux when exposed to KCl nor do they significantly impact the calcium flux of SMA MNs (**C**) without the influence of SMA ACM (t-test,ns). **D**. SMA microglia + MN co-cultures treated with SMA IL1ra/CCL5NAb ACM have significantly higher calcium flux after KCl depolarization than SMA UTX ACM or 48wash ACM (1-way ANOVA, *p<0.05). **E**. Phagocytosis of co-cultures can be quantified using RFP+/GFP+ colocalization in microglia (yellow box) via live imaging paradigm. **F**. Quantification of RFP+/GFP+ phagocytosis after 48-hours of ACM treatment confirms decreased phagocytosis in cultures treated with SMA IL1ra/CCL5NAb ACM or 48wash ACM compared to SMA UTX ACM, and no significant benefit of adding in IL-1ra and CCL5 neutralizing antibodies directly onto the co-culture with SMA UTX ACM (1-way ANOVA, ****p<0.0001, ns). **G**. TUNEL assay to assess MN death in SMA microglia + MN co-cultures (63x objective). **H**. Significant decrease in TUNEL+ MNs when co-cultures are treated with SMA IL1ra/CCL5NAb ACM compared to SMA UTX ACM or no treatment (1-way ANOVA, *p<0.05, ****p<0.0001). Representative images taken with Zeiss confocal microscope using a 63x oil objective.

To translate these *in vitro* experiments *in vivo*, we designed a gene therapy targeted to astrocytes by designing an AAV5-packaged viral vector encompassing GFAP-promoted IL-1ra expression linked to an mCherry reporter protein and miRNA knockdown of CCL5 (Figure 5A, +expAAV). A control AAV5-packaged vector was also designed with GFAP promoter driving mCherry and miRNA scramble construct (Figure 5A, +controlAAV). SMNΔ7 (SMA) mice and wild-type (WT) littermates were given intracerebral ventricular (ICV) injections of 2uL of virus per hemisphere containing >10^13^ GC/mL of either +expAAV or +controlAAV on postnatal day 2 (PND2) and tested daily for survival, weight gain, and motor function through time-to-right assay (Figure 5B). mCherry viral expression was found localized to GFAP+ astrocytes in cortex with no differences in number of mCherry+ astrocytes between +controlAAV and +expAAV groups (Figure 5C, 5D, 2-way ANOVA, ns). Expectedly, this was not associated with an increase in SMI-32+ neurons in cortex (Figure 5E, 2-way ANOVA, ns). mCherry expression was also found localized to GFAP+ astrocytes in the lumbar spinal cord (Figure 5F, 5G, 2-way ANOVA, +controlAAV vs +expAAV, ns) where MN loss is known to occur in untreated SMA pups. Importantly, SMA +expAAV mice retained higher numbers of SMI-32+ motor neurons in the lumbar spinal cord than SMA +controlAAV mice (Figure 5H, 2-way ANOVA, *p=0.0272). Both WT and SMA pups +expAAV had approximately 1.5-2-fold increases in IL-1ra protein expression in brain and spinal cord compared to untreated and +controlAAV mice (Figure 6A, 6B, 2-way ANOVAs). SMA +expAAV mice also had a trend for decreased CCL5 expression in spinal cord (Figure 6C, 2-way ANOVA, p=0.1533; Supplemental Figure S3). SMA +expAAV pups had a median survival of 13 days compared to SMA +controlAAV with a median survival of 10.5 days (Figure 6D, 6E, log-rank Mantel-Cox test, **p=0.0049). A modest but significant improvement on motor function was also found, specifically after PND6 (Figure 6F, 2-way ANOVA, SMA +controlAAV vs SMA +expAAV PND7 *p= 0.0371, PND10 *p= 0.0223). During this window, SMA +expAAV pups also had significantly higher weight gain than SMA +controlAAV (Figure 6G, t-test, *p=0.0482). Western blot analyses of endpoint tissues revealed significantly lower expression of phNFkB in cortex though not in spinal cord of SMA +expAAV compared to SMA +controlAAV (Figure 6H, 6I, 2-way ANOVAs, **p=0.0048). TGFb and GFAP expression were not significantly different between +expAAV and +controlAAV groups in these endpoint tissues (Figure 6J, 6K, 2-way ANOVAs, ns). Altogether, these *in vivo* data support the idea of astrocyte-targeted therapeutics to prevent glial-driven motor neuron loss in SMA.

**Figure 5.**
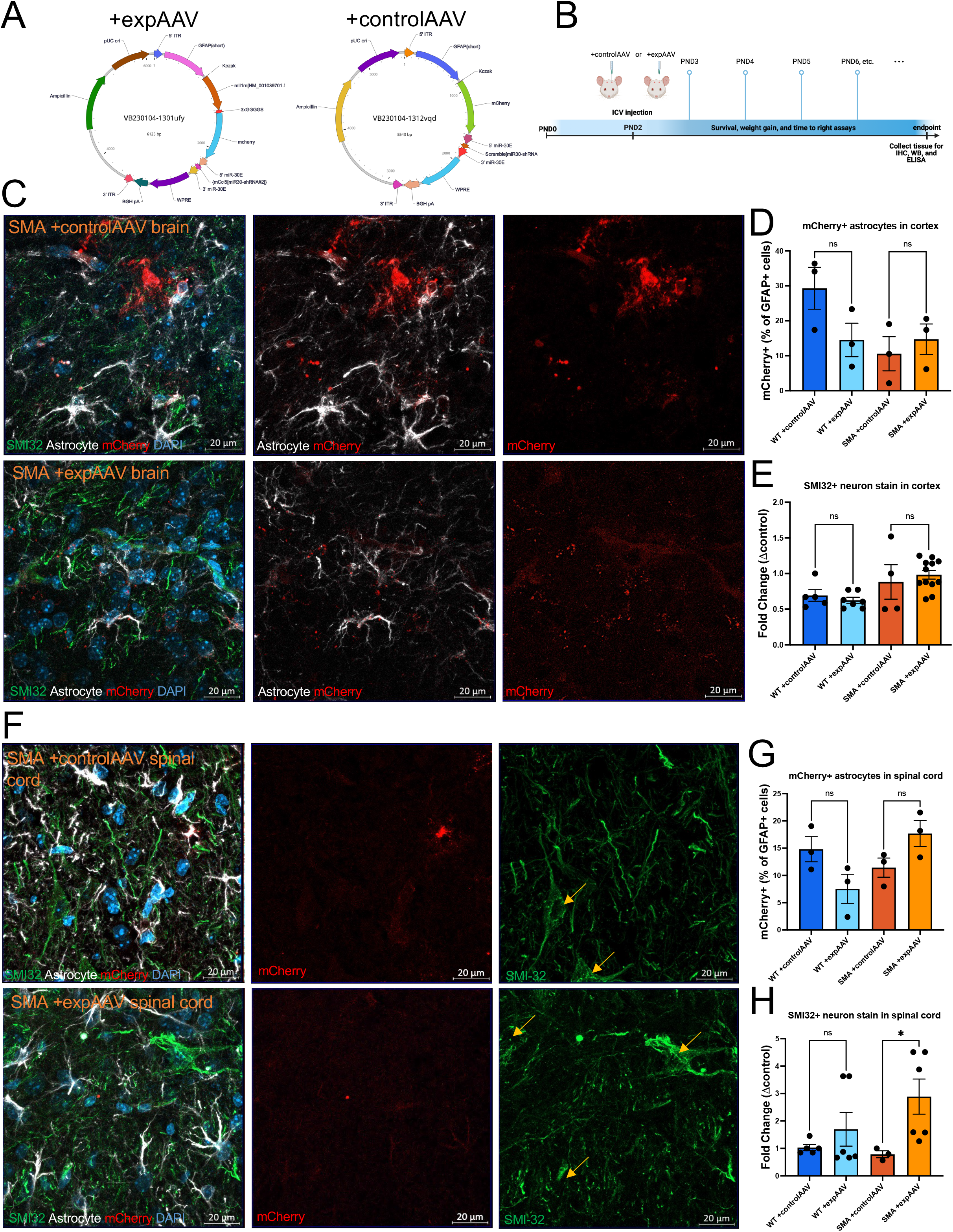
AAV gene therapy to examine effects of IL-1ra increase and CCL5 knockdown in SMNΔ7 mouse model. **A.** Experimental (left, +expAAV) and control (right, +controlAAV) vectors were packaged in AAV5 virions to preferentially target treatment to astrocytes. **B**. 2uL of experimental or control virus (>10^13^ GC/mL) were injected per hemisphere via intracranial ventricular route on PND2. Pups were assessed daily for survival, weight gain, and motor function via time-to-right assay until humane endpoint). **C**. WT and SMA pups show mCherry viral expression localized to GFAP+ astrocytes in cortex (**D**, 2-way ANOVA, ns) and no increase in SMI-32+ MN count between +controlAAV and +expAAV (**E**, 2-way ANOVA, ns). **F**. mCherry+/GFAP+ astrocytes were also found in lumbar spinal cord of +controlAAV and +expAAV WT and SMA pups (**G**, 2-way ANOVA, ns) **H**. SMA +expAAV maintained significantly higher number of SMI-32+ MNs in spinal cord compared to SMA +controlAAV (2-way ANOVA, *p<0.05). Representative images taken with Zeiss confocal microscope using a 63x oil objective. Yellow arrows point out representative SMI-32+ MNs with soma in field of view.

**Figure 6.**
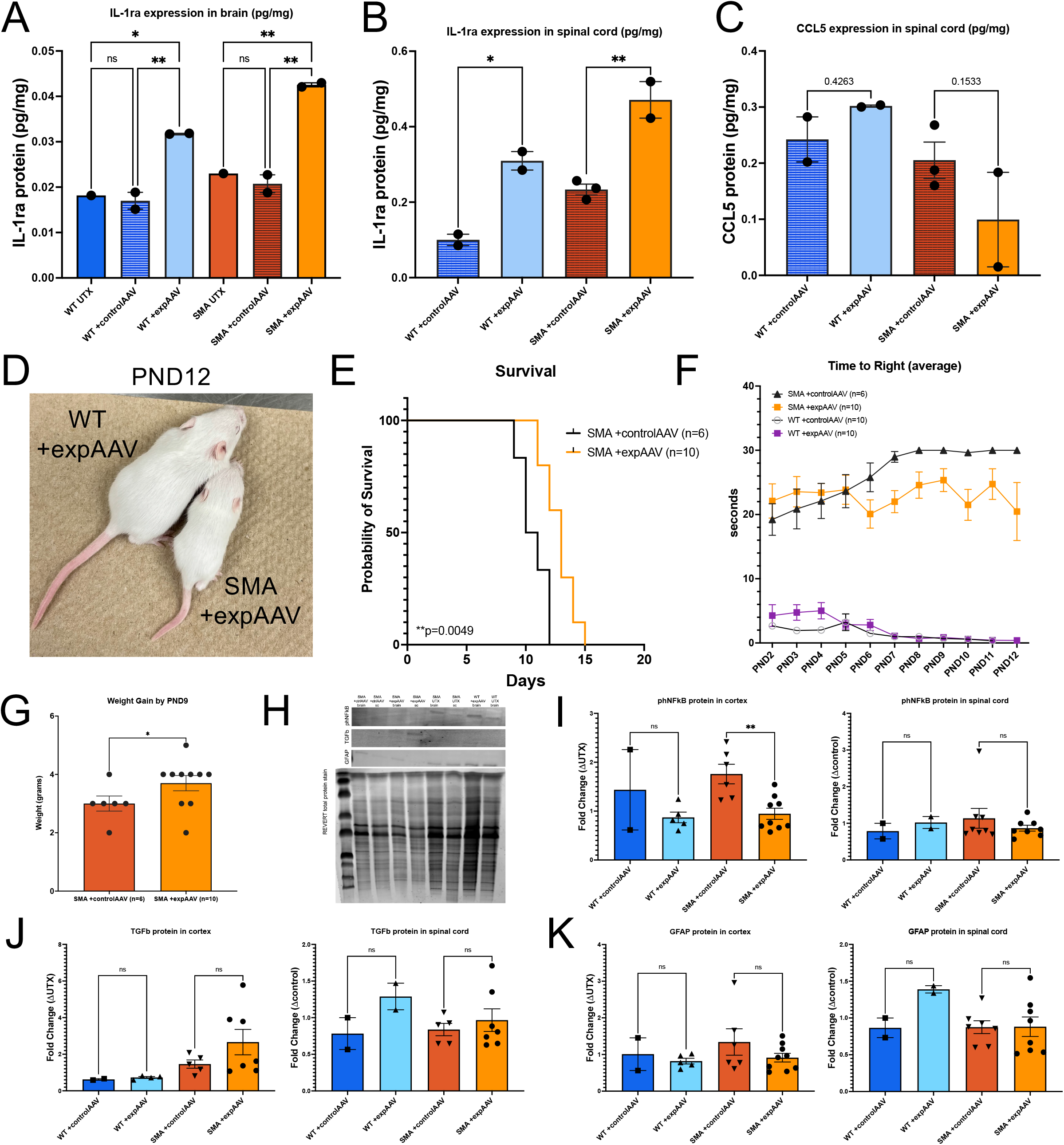
Assessment of IL-1ra and CCL5 treatment on SMA mouse survival and disease phenotypes. WT and SMA pups with +expAAV expressed 1.5-2-fold higher levels of IL-1ra in cortex (**A**) and spinal cord (**B**) than untreated or +controlAAV pups (2-way ANOVA, *p<0.05, **p<0.005). **C**. Trend for lower CCL5 in spinal cord in SMA +expAAV compared to SMA +controlAAV, though no change in WT pups (2-way ANOVA, ns). SMA +expAAV (**D**, representative image next to WT littermate) survived longer than SMA +controlAAV pups (**E**, Mantel-Cox test, **p<0.005). **F**. Modest effect on SMA +expAAV motor function was also seen compared to SMA +controlAAV during PND6-PND12 window (2-way ANOVA, *p<0.05 on PND7 and PND10). **G**. Weight gain was also significantly increased during this window in SMA +expAAV compared to SMA +controlAAV (t-test, *p<0.05). **H**. Western blot analyses at endpoint reveal a significant reduction in phNFkB protein in cortex of SMA +expAAV compared to SMA +controlAAV pups though no significant reduction in spinal cord (**I**, 2-way ANOVAs, **p<0.005, ns). **J**. TGFb protein had a trend for increased expression in cortex, though no significant differences in spinal cord (2-way ANOVAs, ns). **K**. No significant differences in GFAP expression in brain or spinal cord with +expAAV (2-way ANOVAs, ns).

Finally, we assessed the levels of CCL5 and IL-1ra in SMA patient cerebrospinal fluid (CSF) to determine if these pathways may be altered by increasing levels of SMN or associated with improved patient phenotypes. CSF was collected from SMA patients before receiving injections of the antisense oligonucleotide (ASO) treatment nusinersen (Table 1). SMA type 1 patients had high levels of CCL5, which did not decrease over the course of treatment (Figure 7A, repeated measures 2-way ANOVAs with Geisser-Greenhouse corrections, ns). One Type 2 patient (Figure 7B, PT 18) and one Type 3 patient (Figure 7C, PT 23) had a small but significant reduction in CCL5 after more than 2 ASO treatments (PT 18 *p=0.0333, PT 23 *p=0.0182). Patients who were genetically confirmed during newborn screening to have SMA but received treatments before the onset of any symptoms (pre-symptomatic) also did not have any reductions in CCL5 over the course of treatment (Figure 7D, ns). Both SMA type 1 patients were found to have reduced IL-1ra levels from pre-treatment to <2 ASO treatments (Figure 7E, PT 6 ****p<0.0001, PT 20 *p=0.0336). Interestingly, only the SMA type 2 and SMA type 3 patients that did not have decreased CCL5 were found to have increased IL-1ra (Figure 7F, PT 10 **p=0.0075, Figure 7G, PT 4 *p=0.0252). Although pre-symptomatic patient 25 had some variation in IL-1ra level (Figure 7H), neither pre-symptomatic patient had significantly different IL-1ra levels from pre-treatment to >4 ASO treatments (PT 25 pre vs >4 ns, PT 28 pre vs >4 ns). These data indicate that increasing SMN is not sufficient to reduce pro-inflammatory glial phenotypes and provide support for an astrocyte-targeted therapeutic to be considered in combination with FDA-approved therapies to restore SMN.

**Figure 7.**
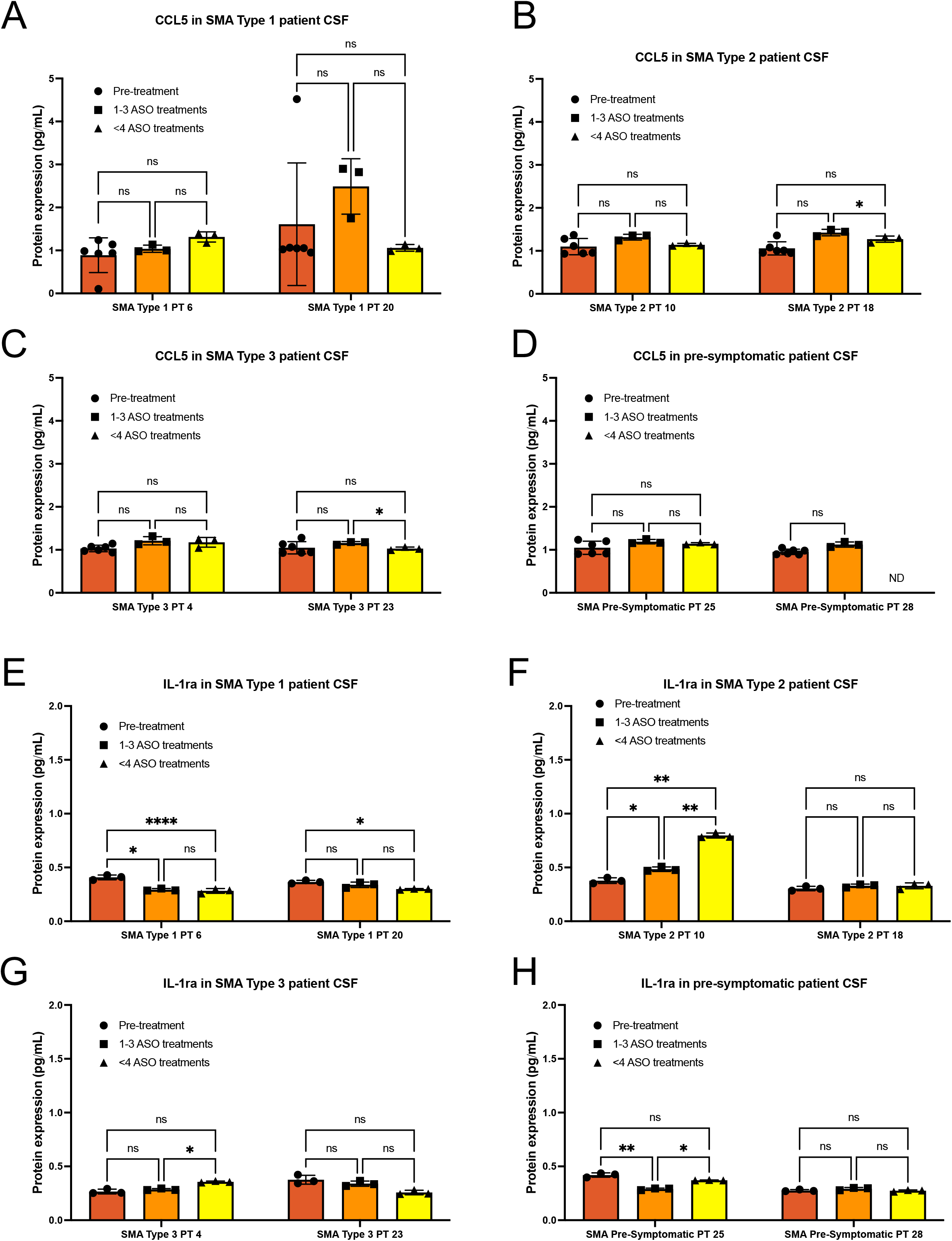
Levels of CCL5 and IL-1ra in SMA patient CSF are not improved with nusinersen treatment. Cerebrospinal fluid (CSF) from SMA patients collected before first antisense oligonucleotide (ASO, nusinersen) treatment (pre-treatment), after 1-3 ASO treatments, and after at least 4 treatments (<4 ASO). **A**. CCL5 levels were highest in SMA type 1 patient CSF and did not significantly decrease over course of treatment. **B**. One SMA type 2 and one SMA type 3 (**C**) had small but significant decreases in CCL5 whereas other patients and pre-symptomatic patients (**D**) did not. E. IL-1ra expression levels decreased in SMA type 1 patients over treatment course, whereas the type 2 (**F**) and type 3 (**G**) patients who did not have decreased CCL5 had increased IL-1ra. **H**. Pre-symptomatic patients had some variation in IL-1ra expression but at latest treatment timepoint were not significantly different from pre-treatment values (repeated measures 2-way ANOVAs with Geisser-Greenhouse corrections, *p<0.05, **p<0.005, ****p<0.0001).

**Table 1.** List of cerebrospinal fluid (CSF) samples from SMA patients.

| SMA Type | Patient | Gender | Age at consent | Pre-treatment sample | 1-3 ASO treatment sample | >4 ASO treatment sample |
| --- | --- | --- | --- | --- | --- | --- |
| 1 | PT 6 | Female | 7 months | CSF001 | CSF003 | CSF004 |
| 1 | PT 20 | Female | 14 years | CSF001 | CSF002 | CSF005 |
| 2 | PT 10 | Male | 13 years | CSF001 | CSF004 | CSF011 |
| 2 | PT 18 | Female | 3 years | CSF001 | CSF004 | CSF010 |
| 3 | PT 4 | Male | 9 years | CSF001 | CSF004 | CSF018 |
| 3 | PT 23 | Male | 4 years | CSF001 | CSF004 | CSF006 |
| Pre-symptomatic | PT 25 | Female | 85 days | CSF001 | CSF003 | CSF004 |
| Pre-symptomatic | PT 28 | Male | 5 years | CSF001 | CSF003 | CSF004 |
| Mean |  |  | 6.1 years | 1 | 3.5 | 5.5 |

## Discussion

SMA patient clinical data has found that early, pre-symptomatic SMN treatment is the most effective for improving survival, motor function, and clinical milestone scores in treated patients^48^. Similarly, mouse models of severe SMA have shown that a short window exists in which widespread AAV9-mediated restoration of SMN is most beneficial^23^, such that therapeutic efficacy is diminished when delivered after PND5^21^. Because early astrocyte malfunction is observed prior to motor neuron loss^7,49^, it is likely that these neuron-targeted interventions do not ameliorate astrocyte pathology including upregulated production of pro-inflammatory cytokines^7-10^ General anti-inflammatory treatments have been proposed for SMA patients after promising results were seen in mouse models^50-52^; however, these treatments have not shown enough benefit to SMA patients to make it through clinical trials^52^. This indicates that a general anti-inflammatory approach is not sufficient and provides support for a disease-specific treatment model targeting astrocyte-driven inflammation for SMA.

Here, we tested IL-10 as a therapeutic target *in vitro* and combined this anti-inflammatory treatment with a neutralizing antibody treatment against CCL5 – the most upregulated pro-inflammatory cytokine secreted by SMA astrocytes (Figure 1). We did not find IL-10 capable of reducing the pro-inflammatory phenotype of SMA astrocytes, even when astrocytes were treated with twice the amount of IL-10 found in HC microglia capable of reducing astrocyte-driven phenotypes in other disease contexts^37^. IL-10 receptor activity has been shown to be reduced in aged astrocytes with aberrant activation phenotypes^53^, so it is possible that SMA astrocytes also have decreased activity of this receptor that is not linked to expression levels of IL-10. We did find decreased CCR5 signaling in IL10/CCL5NAb-treated astrocytes and minor effects of IL10/CCL5NAb ACM on microglial phenotypes to indicate a potential benefit of CCL5 neutralization (Figure 2). The only anti-inflammatory phenotype we were able to identify in IL10/CCL5NAb-treated astrocytes was an increase in secretion of IL-1ra, which only occurred when astrocytes were given a continuous 96-hour treatment (Figure 3). NFkB can be inhibited by reduced CCR5 signaling via increased pCREB^54^, so this extended CCL5 neutralization may have allowed for an increase in this anti-inflammatory protein. IL-1ra acts via binding of the IL-1 receptor to prevent signaling of IL-1β and IL-1a^35^. SMA astrocytes do produce high levels of IL-1β9,10, which can contribute to propagated pro-inflammatory signaling through autocrine stimulation of IL-1R and downstream NFkB activation^55^. Recombinant IL-1ra is already approved to treat rheumatoid arthritis and several other inflammatory diseases^56,57^ and so was a promising replacement for IL-10 in our SMA astrocyte-targeted treatment paradigm.

IL-1ra treatment combined with CCL5 neutralization was successful at reducing SMA astrocyte pro-inflammatory transcript and protein levels, decreasing SMA astrocyte-driven microglial priming and phagocytosis, and decreasing astrocyte-induced MN dysfunction *in vitro* (Figures 3, 4). Unlike co-cultures of SMA motor neurons with SMA astrocytes^38^, direct contact with SMA microglia did not affect MN function or survival without the influence of SMA astrocyte conditioned media (Figure 4). This supports the hypothesis that SMA microglia are activated by SMA astrocyte secreted factors like CCL5. Consistent with previous studies which found microglia to respond better to astrocyte secreted factors than direct treatments^58,59^, direct treatment of SMA microglia did not have the same effect as IL1raCCL5 treated astrocyte media on reducing phagocytosis (Figure 4F). Additionally, we have previously found no effect of healthy microglia conditioned media on SMA astrocyte phenotypes^10^, so microglia-directed treatments would likely not correct this astrocyte-driven inflammation.

Because of the success of IL-1ra and CCL5 neutralization on reducing astrocyte-driven pathology *in vitro*, we next designed an AAV5-based gene therapy to test the effects of these astrocyte-targeted therapeutics *in vivo* using an SMNΔ7 mouse model (Figure 5, 6). We hypothesized that astrocyte-targeted overexpression of IL-1ra and miR-based short hairpin RNA knockdown (miR30-shRNA) of CCL5 would prevent astrocyte-driven microglial activation and motor neuron loss. We predicted that this treatment would delay the onset of motor symptoms and reduce glial activation, but without correcting the SMN deficiency would not reverse SMA phenotypes. We found expression localized to astrocytes throughout the brain and spinal cord of injected mice (Figure 5), though expression was not readily apparent in all GFAP+ astrocytes. IL-1ra expression was increased 1.5-2-fold in treated animals and associated with decreased phNFkB expression in cortex (Figure 6); however, CCL5 expression did not appear significantly reduced. shRNA constructs often do not have complete knockout effects, with 50-70% having noticeable knockdown and only 20-30% having a strong knockdown effect^60,61^. Future experiments may examine other CCL5 shRNA constructs or alternative knockdown strategies. One such strategy could include overexpression of miR-324-5p, a microRNA that is significantly decreased in SMA type 1 patient serum^31,62^. Treatment of SMA mice with miR-324-5p has been found to improve weight gain for a short period^31^ and may be a promising alternative to a direct CCL5 knockdown.

Despite our IL1a/CCL5NAb therapy having modest overall improvements on lifespan, weight gain, and motor function, analyses of endpoint tissues (PND ∼10.5 for SMA +control AAV, PND ∼13 for SMA +expAAV) did not reveal significant changes to TGF-β, GFAP, or phNFkB expression in the spinal cord (Figure 6). It will be important to collect tissues along the time course of treatment to directly compare the onset of inflammation in control injected vs treated SMA mice. It is also possible that the astrocytes not expressing the transgene are still contributing to pro-inflammatory state and motor neuron loss, so increasing viral titer or efficiency may help correct these issues. Finally, SMA patients were overall not found to have consistent decreases in CCL5 or increases in IL-1ra after treatment with nusinersen, the ASO therapeutic that modulates the pre-mRNA splicing of *SMN2* to create more full length SMN protein^63,64^ (Figure 7). These individual patient data fit with the recent finding that upregulated pro-inflammatory cytokines in SMA patient CSF compared to controls were not decreased after nusinersen treatment^65^. To directly examine the effects of IL-1ra/CCL5NAb treatment with SMN restoration, future studies will include treating +expAAV animals daily with nusinersen. These studies will provide valuable insight into the effect of combination therapy of nusinersen with IL-1ra on pro-inflammatory status and patient outcomes in more severe SMA patient types.

Together, these data provide insight into the mechanisms of astrocyte-driven inflammation and disease pathology in SMA. We specifically identify IL-1ra and CCL5 as therapeutic targets that can be translated to combinatorial treatments to improve patient function and quality of life. Finally, these experiments highlight the importance of cell- and disease-specific therapeutic design.

## Materials and methods

### Cell culture

Two healthy control (21.8, 4.2) and three SMA patient (7.12, 3.6, 8.2) iPSC lines were utilized in these experiments^8,10^. All pluripotent stem cells were maintained on Matrigel (Corning) in Essential 8 (Gibco) and passaged every 4-6 days. iPSCs and differentiated cells were confirmed mycoplasma negative. SMA 7.12 and SMA 8.2 were made into GFP- and RFP-expressing stable lines by infecting cells with LentiBrite GFP Control Lentiviral Biosensor (Millipore, #17-10387, titer 7.34 x108 IFU/mL) or LentiBrite RFP Control Lentiviral Biosensor (Millipore, #17-10409, titer 4.59 x 108 IFU/mL) at MOI of 20 for 24 hours. Virus was removed and cells were allowed to expand for 1 week. GFP+ and RFP+ cells were isolated using WOLF Cell Sorter (Nanocellect) and expanded to create purified stable lines.

### Astrocyte, microglia, and motor neuron differentiations

Spinal cord patterned astrocytes were generated from iPSCs as previously described^10,37^. Differentiation reagents were purchased from ThermoFisher unless otherwise noted. Briefly, iPSCs were grown to confluency, dissociated with Accutase, and differentiated towards NPCs. NPCs were passaged via Accutase treatment every 6 days. P3 NPCs were used for astrocyte differentiations and cultured in ScienCell Astrocyte Medium (ScienCell Research Laboratories, Carlsbad, CA, USA) containing 1% astrocyte growth supplement, 1% penicillin– streptomycin, and 2% B27. Cells were fed every 48h and passaged with Accutase every 6–9 days upon confluency (minimum of 3 passages). Passage 4 cells were considered fully differentiated and seeded onto T75 tissue culture flasks for ACM generation and collection.

Microglia were differentiated using the commercially available differentiation kit (StemCell Technologies #05310, #100-0019, #100-0020, Vancouver, BC, Canada) based on a previously published protocol^66^. As previously described^10,37^, iPSCs were differentiated into hematopoietic progenitor cells (HPCs) using the STEMdiff Hematopoietic Kit (StemCell Technologies, Vancouver, BC, Canada). Floating HPCs were collected and plated at 50,000 cells/mL in STEMdiff microglia differentiation media (StemCell Technologies, Vancouver, BC, Canada) for 24 days, followed by rapid maturation in STEMdiff microglia maturation media (Stem-Cell Technologies, Vancouver, BC, Canada) for a minimum 4 days.

Spinal motor neurons were differentiated based on the Maury et al. (2015) protocol^67^. Briefly, embryoid bodies were generated from iPSCs and patterned in the presence of Chir-99021 with dual SMAD inhibition (SB 431542 and LDN 1931899) followed by treatment with retinoic acid (RA), smoothened agonist (SAG), and DAPT. Spinal motor neuron progenitor cells were then dissociated and plated on Matrigel-coated glass coverslips or 96-well plates for terminal differentiation and maturation in growth factor supplemented medium for 21–42 days *in vitro*.

### Astrocyte immunocytochemistry

Plated astrocytes were fixed in 4% paraformaldehyde (PFA) for 20 min at room temperature and rinsed with PBS. Nonspecific labeling was blocked and the cells permeabilized with 0.25% Triton X-100 in PBS with 1% BSA and 0.1% Tween 20 for 15 minutes at room temperature. Cells were incubated with primary antibodies overnight at 4C, then labeled with appropriate fluorescently-tagged secondary antibodies. Hoechst nuclear dye was used to label nuclei. The primary antibody used was rabbit anti-GFAP (GFAP; DAKO #Z0334) at 1:500 dilution with secondary antibody donkey anti-rabbit AF546 (Invitrogen, Waltham, MA, USA) used at 1:1000 dilution. Image in Figure 1A was acquired using Zeiss confocal microscope with 63x oil objective and are displayed as a maximum intensity projection of z-stack image series.

### Multiplex human cytokine array on conditioned media

Eve Technologies (Calgary, AB, Canada) performed the 48 multiplex cytokine array from duplicate differentiations using conditioned medium samples generated from iPSC-derived astrocytes. Data were analyzed for fold change difference of each cytokine expression level in SMA ACM compared to HC ACM.

### Astrocyte conditioned media treatments of microglia and MNs

Astrocyte-conditioned media (ACM) were collected upon each media change after P4, spun to remove any cells or debris, and stored in sterile Falcon tubes at −20C. Frozen media were slowly thawed on ice before use. Treatments of MNs with ACM were performed with 25% ACM in MN Maturation media after day 28 and left for 48 hours at 37C before fixing, collection, or analyses. Microglia were treated with 1:2 astrocyte conditioned media with microglia maturation media for 24 hours at 37C. Direct IL10CCL5 or IL1raCCL5 treatments of microglia were given concurrently with SMA UTX ACM treatment and at the same concentration as astrocyte-targeted treatments.

### Microglia soma size and pHrodo phagocytosis assays

Microglia were treated with microglia maturation media (UTX), 1:2 ACM, or 100ng/mL lipopolysaccharide (LPS, Sigma Aldrich, L2018) and placed in Incucyte (Sartorius) to allow for live cell imaging with 20x objective during 24-hour treatment. Soma size was analyzed using Incucyte software to calculate average soma area for all cells in one well (5,000 microglia per well of 24-well plate) at 24-hour timepoint. Points in soma size graphs represent the average of 3 technical well replicates. After 24-hour ACM treatment and soma measurement, pHrodo Red Zymosan Bioparticles (ThermoFisher, #P35364) were added a 1ug/mL to microglia cultures. Plates were returned to Incucyte and imaged for 24 hours using brightfield and red fluorescent channels at 20x. Images were analyzed using Incucyte software for total number of RFP+ microglia in each well and averaged across three technical replicates. Data points on graph represent mean and standard error of the mean for experimental replicates across two separate SMA lines.

### Fluo-4NW calcium flux assay

Calcium flux was measured in MNs seeded in a 96 well plate using the Fluo-4 NW Calcium Assay Kit (ThermoFisher, #F36206) per the manufacturer’s instructions. Growth medium was removed, and 100μL of dye loading solution was added to each well and incubated for 30 minutes at 37C followed by 30 minutes at room temperature. Just before measuring fluorescence, 25μL of the appropriate agonist or PBS (control) were spiked into each well. Fluorescence (excitation 494nm, emission 516nm) was immediately measured using a GloMax microplate reader. No differences in fluorescence were found between pre-agonist background fluorescence and PBS-stimulated wells (Supplemental Figure S1); relative fluorescence (% above PBS baseline) was calculated by subtracting the fluorescence of PBS-stimulated wells from each test well then dividing by PBS-stimulated well value and multiplying by 100. Agonist solutions were made fresh each day and included 250mM KCl (final concentration 50mM), 500μM L-glutamate (final concentration 100μM). Individual data points in relative fluorescence graphs represent the average of three technical replicates.

### *In vitro* astrocyte treatments

Treatments of SMA astrocytes were performed with 430pg/mL recombinant IL-10 (PeproTech, #200-10), 200pg/mL recombinant IL-1ra (PeproTech, #200-01RA), and 20pg/mL CCL5 neutralizing antibodies (Bio-techne, #MAB278-100) in supplemented Astrocyte media for 48 hours before collection of treatment media and cell pellets. For 48 wash conditions, astrocytes were rinsed with PBS and fed with supplemented Astrocyte media. Media were collected from 48wash astrocytes after 48 hours. Treatment values were set based on doubling the amount of IL-10 and IL-1ra secreted by HC astrocytes and doubling the amount of CCL5 secreted by SMA astrocytes as determined by multiplex cytokine array.

### Quantitative real-time polymerase chain reaction (qRT-PCR)

RNA was isolated from cell pellets using the RNeasy Mini Kit (Qiagen) following manufacturer’s instructions, quantified using a Nanodrop Spectrophotometer, treated with RQ1 Rnase-free Dnase (Promega), and converted to cDNA using the Promega Reverse Transcription system (Promega). SYBR green RT-qPCR was performed in triplicate using cDNA and run on the Bio-Rad CFX384 real time thermocycler. Primer sequences shown in Supplemental Table S1. Cq values for each target were normalized to GAPDH and calculated using the ΔΔCq method. A minimum of three differentiations for each line were collected and run in technical triplicates.

### Western blot

Cell pellets and mouse tissues were lysed by sonication with Triton X-100 and then the protein concentration was determined using a BCA assay (ThermoFisher). Equal amounts of protein were loaded onto 10% or 12% pre-cast Tris-HCl Mini-PROTEAN gels (Bio-Rad, Hercules, CA, USA) and proteins separated by electrophoresis, then transferred to PVDF membranes (Bio-Rad). Membranes were blocked for 1 h in Odyssey TBS blocking buffer (LI-COR), followed by overnight primary antibody incubation and 30 min secondary antibody incubation. Quantification was performed with FIJI (ImageJ, National Institutes of Health, Bethesda, MD, USA) and normalized to REVERT total protein stain (LI-COR). Primary antibodies used were rabbit anti IL-10 (abcam, ab133575, 1:1000), SOCS3 (ThermoFisher, #SOCS3-301-AP, 1:500), phSTAT3 (Cell Signaling, #9138S, 1:500), pCREB (ThermoFisher, #MA1-114, 1:500), mouse anti-NFkB p65 (Cell Signaling, #6956, 1:1000) rabbit anti-phNFkB (Cell Signaling, #3033, 1:1000), mouse anti-TGFβ (ThermoFisher, #MA5-16049, 1:1000), rabbit anti-GFAP (DAKO, #Z0334, 1:2000), and mouse anti-C3 (ThermoFisher, #PA5-21349, 1:2000). Secondary antibodies used were anti-rabbit IRDye 800CW (LI-COR, 1:5000 dilution, Lincoln, NE, USA) and anti-mouse IRDye 680RD (LI-COR, 1:5000 dilution, Lincoln, NE, USA).

### Microglia and MN co-cultures

After a minimum of 4 days in STEMdiff maturation media, SMA microglia were added at a ratio of 1:4 to SMA MNs (between 21-36 days of MN maturation) with either 25% SMA UTX, IL1raCCL5, or 48wash ACM, direct IL1raCCL5 treatment with 25% SMA UTX ACM, or 25% fresh Astrocyte media (UTX). Treated co-cultures were placed in Incucyte for 48 hours and imaged every 2 hours in brightfield, RFP, and GFP channels at 20x. After 48 hours, co-cultures were fixed for TUNEL assay or analyzed for calcium flux using Fluo4-NW assay. Phagocytosis was calculated using Incucyte software to identify the number of GFP+/RFP+ double positive microglia at the 48-hour timepoint. Data points on co-culture phagocytosis graph represent the average of three technical replicates for double positive GFP+/RFP+ microglia normalized to total number of cells in the field of view.

### TUNEL assay

Plated cells were fixed in 4% PFA for 20 min at room temperature, rinsed with PBS, and then stained using a Click-iT TUNEL Alexa Fluor 647 Assay Kit (ThermoFisher, #C10247) following manufacturer’s instructions. Briefly, cells were permeabilized as described for ICC, washed, and incubated with DNA labeling solution for 1 hour at 37C. Cells were washed and optional DAPI nuclear counterstain was then applied for 30 min at room temperature. Coverslips were imaged with standard fluorescent microscopy. Three images were acquired from randomly selected fields for each coverslip. Images were analyzed for total fluorescence in either channel using FIJI (ImageJ) software. Relative expression for each condition (total TUNEL stain divided by total 7AAD stain) was analyzed to account for variable number of cells in each ROI as previous described^10^. Representative images were acquired on a Zeiss confocal microscope using 63x oil objective and are displayed as a maximum intensity projection of z-stack image series.

### *In vivo* AAV vector design and injections

Plasmids were designed using VectorBuilder. Experimental virus (+expAAV) was designed with a GFAP (short) promoter driving expression of mouse IL-1ra (NM_001039701.3) linked to mCherry reporter protein via 3xGGGGS and miR30-shRNA knockdown of mouse CCL5. Control virus (+controlAAV) was designed with GFAP (short) promoter driving expression of mCherry and contained an miR30-shRNA scramble construct. VectorBuilder prepared the plasmid and performed the AAV5 packaging and purification. Final control and experimental viruses contained >10^13^ GC/mL.

### Animals

Mice were housed and handled in accordance with the Animal Care and Use Committee of the Medical College of Wisconsin. SMNΔ7 mice were purchased form JAX Laboratories (stock numbers xxx). SMNΔ7 homozygous mutants (SMA mice) had a homozygous deletion of the mouse Smn gene and homozygous transgenes for human SMN2 and an SMN cDNA lacking exon 7. Untreated SMA mice live to ∼12 days and show signs of progressive muscle weakness, failure to thrive (indicated by weight gain), and a loss of lower motor neurons in the lumbar spinal cord. SMA mouse pups and unaffected littermates of both sexes received 2uL of +controlAAV or +expAAV per hemisphere on postnatal day 2 (PND2) via intracranial ventricular injections. Mice were assessed daily for survival, weight in grams, and motor function through standard time-to-right assay. Cortex and lumbar spinal cords were collected at humane endpoint for SMNΔ7 mice and matched timepoints for unaffected littermates. Tail biopsies were used for genotyping after injection.

### Immunohistochemistry

10 micrometer sections through cortex and lumbar spinal cords were used for immunohistochemistry analyses. Sections were permeabilized in 0.5% triton X-100 and blocked in 1% BSA. Sections were incubated in primary antibody overnight at 4C and incubated with secondary antibodies for 1 hour at room temperature. Primary antibodies used were mouse anti-SMI-32 (Covance, #SMI-32R, 1:500), rabbit anti-GFAP (DAKO, #Z0334, 1:500), and mCherry Monoclonal Antibody (16D7) AF 594 (ThermoFisher, #M11240, 1:500). Secondary antibodies used were donkey anti-mouse AF488 (Invitrogen) and donkey anti-rabbit AF647 (Invitrogen). Analyses were performed on a minimum of three randomly selected fields per condition using standard fluorescent microscopy and equivalent exposure conditions at 63x. Images were analyzed for total fluorescence in each channel using Nikon Elements software. mCherry+ astrocytes were calculated using the Object Count function for mCherry only channel and represented as a percentage of GFAP+ objects in the GFAP only channel. SMI-32+ neuron stain is represented as the fold change of total SMI-32 signal from a randomly selected +controlAAV image for WT and SMA mice. Representative images were acquired using a confocal microscope with a 63x oil objective and were displayed as the maximum intensity projection (MIP) of a z-stack of images.

### Mouse tissue IL-1ra and CCL5 ELISAs

Cortex and lumbar spinal cord sections were weighed (in mg) before lysis by manual pipetting and sonication with Triton X-100, and then the protein concentration was determined using a BCA assay (ThermoFisher). Samples were analyzed using RANTES Mouse Instant ELISA Kit (Invitrogen, #BMS6009INST) and IL-1RA Mouse ELISA Kit (Invitrogen, #EMIL1RN). Calculated protein concentrations were then normalized to starting tissue weight to generate values in pg/mg.

### Human samples

CSF samples were obtained at Children’s Wisconsin with informed consent during standard clinical care for the administration of SMA-related nusinersen ASO therapy (IRB 1080061). A total of 24 CSF samples were used in this study from SMA Type 1 (2 patients), Type 2 (2 patients) and Type 3 (3 patients) as well as 2 pre-symptomatic patients (Table 1). Deidentified SMA patient information related to pre– and post–nusinersen-treated CSF biospecimens analyzed in this study are detailed in Table 1. Before each dose, the patients received a physical exam by a pediatric neuromuscular physician or physician’s assistant as well as complete blood count, urinalysis, complete metabolic panel, INR, PT and PTT as previously described^68^. CSF samples were immediately frozen after collection and stored at −80°C. CSF samples visibly contaminated with blood were not used in this study (PT 25 <4 ASO treatment data for CCL5 was excluded for this reason). The use of CSF samples and iPSCs was approved by the Medical College of Wisconsin Institutional Review Board (PRO00030075; PRO00025822), the Institutional Biosafety Committee (IBC20120742) and the Human Stem Cell Research Oversight Committee. Samples were analyzed using Human RANTES ELISA Kit (ThermoFisher, EHRNTS) and IL-1RA Human ELISA Kit (ThermoFisher, KAC1181). Protein concentrations were calculated per manufacturer’s instructions in pg/mL. For CSF001 samples, duplicate experiments were run in technical triplicates, all other sample data shown are technical triplicates.

### Statistical analyses

Experimental conditions within each experiment were performed in technical triplicates for a minimum of three independent experiments unless otherwise noted. Data were analyzed using GraphPad Prism software and the appropriate statistical tests including the Student’s t test, 1-way ANOVA, and 2-way ANOVA followed by Tukey’s post hoc analysis of significance. Changes were considered statistically significant when p<0.05.

## Data availability statement

For original data, please contact.

## Acknowledgements

We would like to thank and acknowledge patients and families for consenting to CSF collection and use. We would also like to thank Anvitha Sriram for technical assistance with genotyping and Rebecca J. Rehborg for coordinating CSF sample collection. This project was supported by the Medical College of Wisconsin Center for Immunology (RLA) and Advancing a Healthier Wisconsin (ADE). MS was supported through the Summer Program for Undergraduate Research at MCW. Figures 1C, 2A 2B, 3B, 3C and 5B created using BioRender.

## Author Contributions

RLA, MS, and ML performed and analyzed experiments. All authors designed experiments and interpreted data. ADE supervised the study and provided funding. RLA wrote the manuscript and created figures. All authors edited and approved the manuscript.

