## Supplemental tables and figure for "IL-1Ra and CCL5, but not IL-10, are promising targets for treating SMA astrocyte-driven pathology"

Supplemental Figures – IL-1ra and CCL5 to treat SMA astrocyte-driven pathology

Supplemental Table S1 – Primer sequences

| Target Gene | Forward Sequence | Reverse Sequence |
|---|---|---|
| NFkB | GCAGCACTACTTCTTGACCACC | TCTGCTCCTGAGCATTGACGTC |
| IL-1B | CCACAGACCTTCCAGGAGAATG | GTGCAGTTCAGTGATCGTACAGG |
| IL-6 | AGACAGCCACTCACCTCTTCAG | TTCTGCCAGTGCCTCTTTGCTG |
| C1q | CAACACAGGCTGCTACGGGATC | CTGCCCTTTGGGTCCTCGGAT |
| C3 | GTGGAAATCCGAGCCGTTCTCT | GATGGTTACGGTCTGCTGGTGA |
| GDNF | CGCCGAAGACCGCTCCCTCG | ATCCATGACATCATCGAACTGATC |
| BDNF | CATCCGAGGACAAGGTGGCTTG | GCCGAACTTTCTGGTCCTCATC |

Supplemental Figure S1 – Quantification of high throughput Fluo-4 NW calcium flux assay

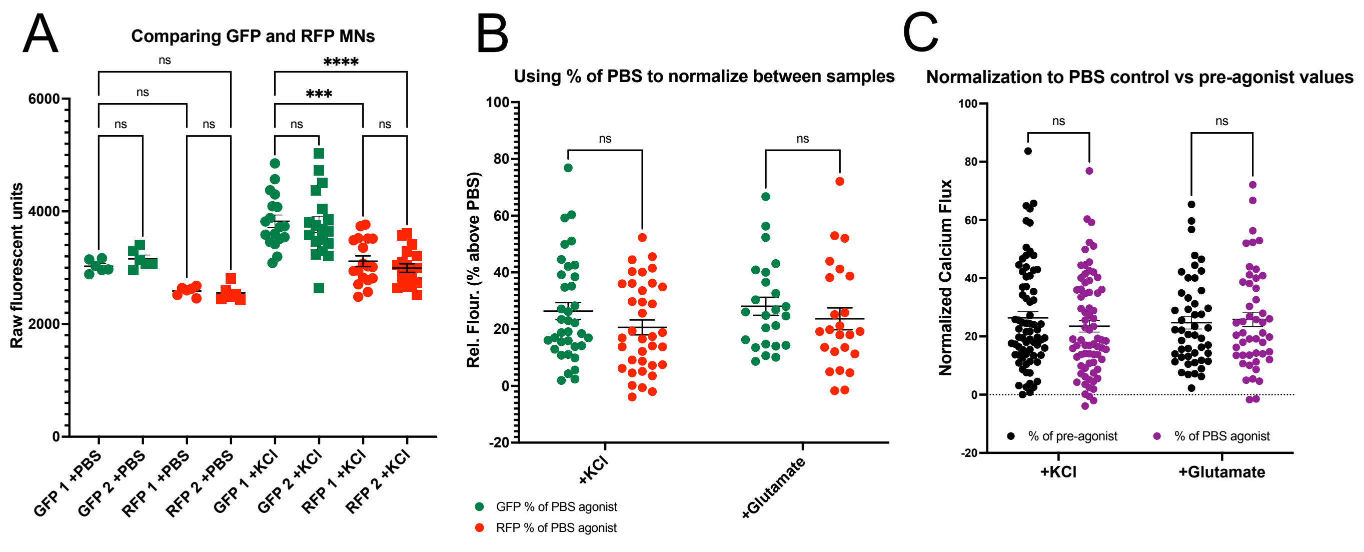


**Supplemental Figure S1 – Quantification of high throughput Fluo-4 NW calcium flux assay. A**. Fluorescence in raw units is higher in GFP-labeled MNs than RFP-MNs due to background fluorescence (1-way ANOVA, ***p<0.0005, ****p<0.0001). **B**. Calcium flux calculated as the percentage of increased florescence in stimulated (+KCl or +glutamate) above unstimulated (PBS added as negative control instead of agonist) wells allows for normalization between GFP- and RFP-labeled MNs. **C**. Using unstimulated wells as a control to normalize % increase in calcium flux (% of PBS agonist) is no different from calculating individual well % increase using a scan taken before agonist is added (% of pre-agonist) (2-way ANOVA, ns).

Supplemental Figure S2 – TUNEL only channel from representative images

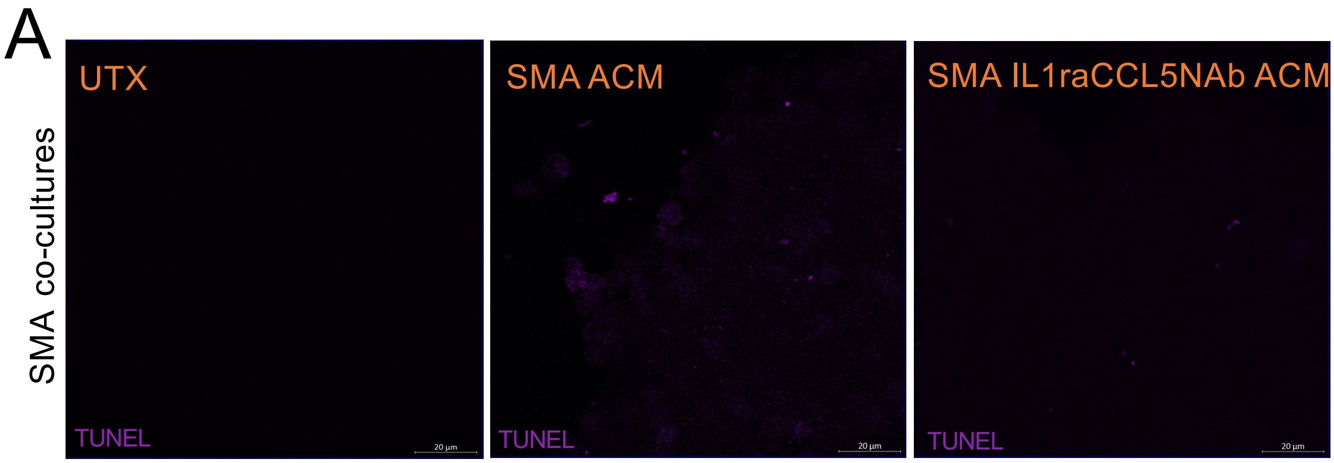


**Supplemental Figure S2 – TUNEL only channel from representative images. A**. TUNEL channel only exports from Figure 4G representative images taken with Zeiss confocal microscope using a 63x oil objective.

Supplemental Figure S3 – Representative images of WT littermates' brain and spinal cord.

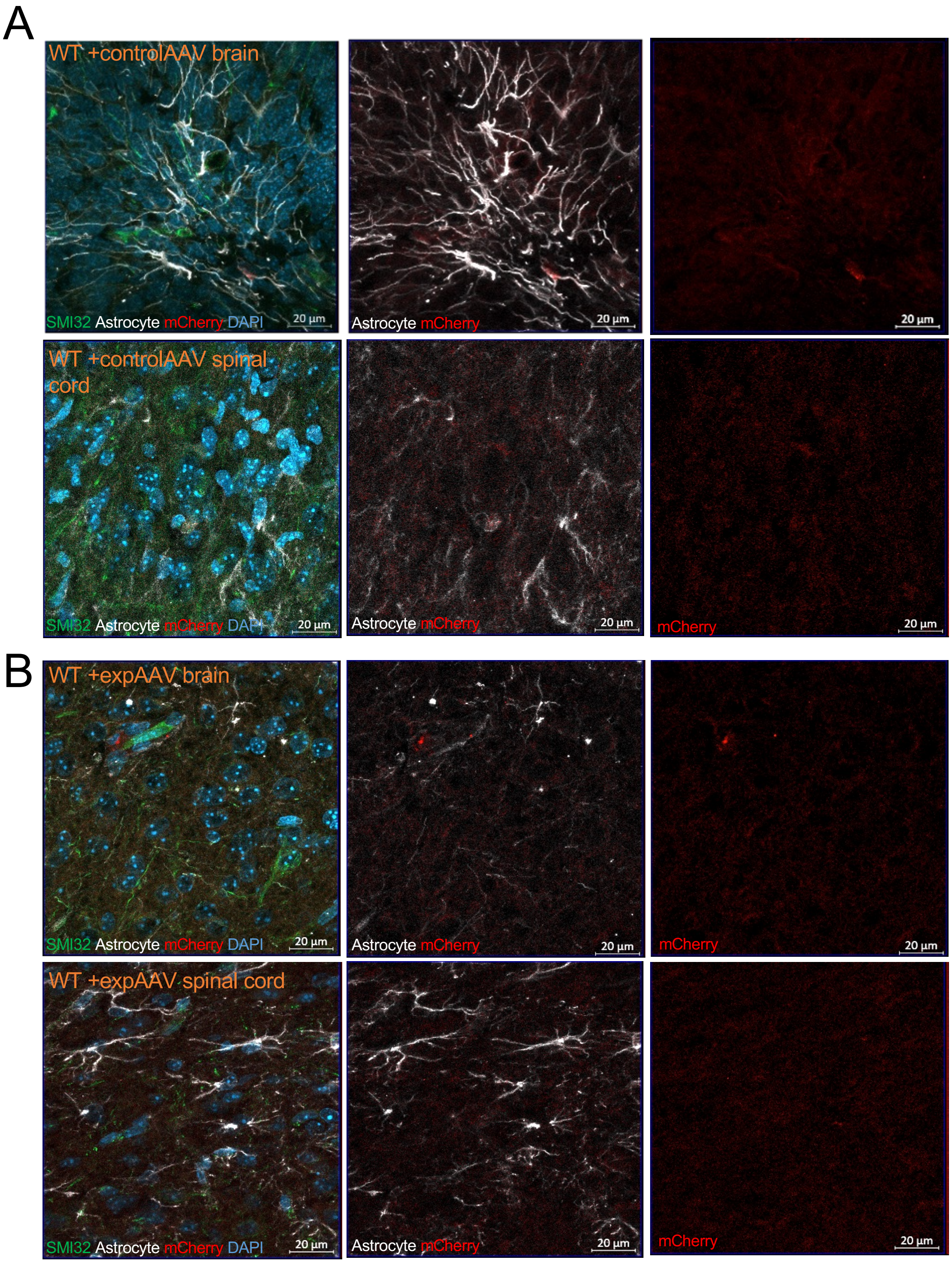


**Supplemental Figure S3 – Representative images of WT littermates' brain and spinal cord.** WT littermates +controlAAV (**A**) or +expAAV (**B**) brain (top) and spinal cord (bottom) stained for SMI-32 (green), GFAP (white), and mCherry viral proteins (red). Representative images taken with Zeiss confocal microscope using a 63x oil objective.
